# WGDI: A user-friendly toolkit for evolutionary analyses of whole-genome duplications and ancestral karyotypes

**DOI:** 10.1101/2021.04.29.441969

**Authors:** Pengchuan Sun, Beibei Jiao, Yongzhi Yang, Lanxing Shan, Ting Li, Xiaonan Li, Zhenxiang Xi, Xiyin Wang, Jianquan Liu

**Author notes:** These authors contributed equally to the work (Pengchuan Sun, Beibei Jiao). Correspondence should be addressed to Jianquan Liu.

## Abstract

Evidence of whole-genome duplications (WGDs) and subsequent karyotype changes has been detected in most major lineages of life on Earth. To clarify the complex resulting multiple-layered patterns of gene collinearity in genome analyses there is a need for convenient and accurate toolkits. To meet this need, we introduce here WGDI (<u>W</u>hole-<u>G</u>enome <u>D</u>uplication <u>I</u>ntegrated analysis), a Python-based command-line tool that facilitates comprehensive analysis of recursive polyploidizations and cross-species genome alignments. WGDI supports three main workflows (polyploid inference, hierarchical inference of genomic homology, and ancestral chromosomal karyotyping) that can improve detection of WGD and characterization of related events. It incorporates a more sensitive and accurate collinearity detection algorithm than previous softwares, and can accelerate WGD-related karyotype research. As a freely available toolkit at GitHub (https://github.com/SunPengChuan/wgdi), WGDI outperforms similar tools in terms of efficiency, flexibility and scalability. In an illustrative example of its application, WGDI convincingly clarified karyotype evolution in *Aquilegia coerulea* and *Vitis vinifera* following WGDs and rejected the hypothesis that *Aquilegia* contributed as a parental lineage to the allopolyploid origin of core dicots.

## Introduction

There is clear evidence that whole-genome duplication (WGD), or polyploidy, and the accompanying change in karyotype, have repeatedly occurred in diverse eukaryotic lineages (Van de Peer, Mizrachi et al. 2017). It is recognized as a prominent evolutionary process, especially in plants (Soltis and Soltis 2016, Landis, Soltis et al. 2018). Thus, identifying WGDs, their dates and locations in evolutionary history, and ancestral karyotypes, is crucial for thorough understanding of how eukaryotes have diversified and adapted to different environments (Fawcett, Maere et al. 2009, Mabry, Brose et al. 2020). To date, three main types of methods have been used to detect WGD: Ks-based, gene tree-based, and synteny-based methods (Rabier, Ta et al. 2014, Mabry, Brose et al. 2020). Previous studies have shown that Ks-based or gene tree-based methods alone can be potentially misleading (Hahn 2007, Vanneste, Van de Peer et al. 2013, Ruprecht, Lohaus et al. 2017, Tiley, Barker et al. 2018, Nakatani and McLysaght 2019, Zwaenepoel, Li et al. 2019). In contrast, synteny-based methods are more reliable, conclusive and thus currently serve as the gold standard for inferring WGD (Kellis, Birren et al. 2004, Nakatani and McLysaght 2019).

With the increasing availability of assembled genomes, a number of methods have been developed to identify conserved syntenic blocks in eukaryotes, i.e. preserved co-localization of homologous genes on chromosomes. Early software packages developed for this purpose, such as ADHoRe (Vandepoele, Saeys et al. 2002) and DiagHunter (Cannon, Kozik et al. 2003), often relied on clustering of neighboring matching gene pairs. In contrast, more recent packages use dynamic programming to build chains of pairwise collinear genes. Examples include ColinearScan (Wang, Shi et al. 2006), Cyntenator (Rödelsperger and Dieterich 2010), McScan (Tang, Wang et al. 2008), MCScanX (Wang, Tang et al. 2012) and JCVI (Tang, Vivek et al. 2017). However, these software packages lack sufficient compatibility with the Windows platform, and the algorithms they incorporate lack sufficient sensitivity to collinearity, and hence accuracy. Due to the high complexity of extant genomes after often recursive WGDs and subsequent genome reconfiguration (often with extensive chromosomal rearrangement and massive gene loss), capacities for detailed downstream analysis are essential, such as generation of homologous gene dotplots or circles displaying collinear genes or blocks, Ks calculations, Ks peak fits, exploration of ancestral karyotype evolution, and comparison of syntenic genes. No previous software or toolkit provided all these capacities and/or the ability to integrate different software, thus hindering WGD research and frequently leading to erroneous interpretation of ancient polyploidization patterns (Wang, Sun et al. 2018, Wang, Yu et al. 2020).

To facilitate WGD analysis, here we introduce a convenient toolkit called WGDI (<u>W</u>hole-<u>G</u>enome <u>D</u>uplication <u>I</u>ntegrated analysis), a convenient Python-based command-line toolkit with the following advantages. It incorporates a more sensitive collinearity algorithm than previous packages, which can improve the resolution and accuracy of synteny block analyses. It also provides integrated capacities for nearly all current WGD-related and bioinformatic analyses, including (among others) inter- and intra-genomic dotplot comparison, collinearity detection, Ks estimation and peak fitting, ancestral karyotype evolution, and inference of synteny trees. Moreover, parameters for all these analyses can be very simply adjusted.

## Results

### Structure of the WGDI package

The complete WGDI source code is freely available at GitHub (https://github.com/SunPengChuan/wgdi) and can be deployed in Windows, Linux, or macOS operating systems. WGDI is written in python3 and can be installed via pip or conda. WGDI supports three main workflows: (1) analysis and inference of polyploidy using homologous dotplots, collinearity and Ks distributions, and homologous gene trees; (2) hierarchical inference of genomic homology resulting from recursive paleopolyploidization; (3) subgenomic and ancestral chromosomal karyotyping and analysis of evolutionary scenarios. WGDI has multiple subroutines, and users only need to simply modify the configuration file and enter their names (e.g., ‘wgdi -d your.conf’) to execute them. The subroutine parameters and functions of WGDI are shown in Fig. 1. WGDI outputs may include vector diagrams (e.g. in SVG format) that are suitable for direct publication. Detailed function descriptions and parameter settings are available at https://wgdi.readthedocs.io/en/latest/usage.html.

**Fig. 1.**
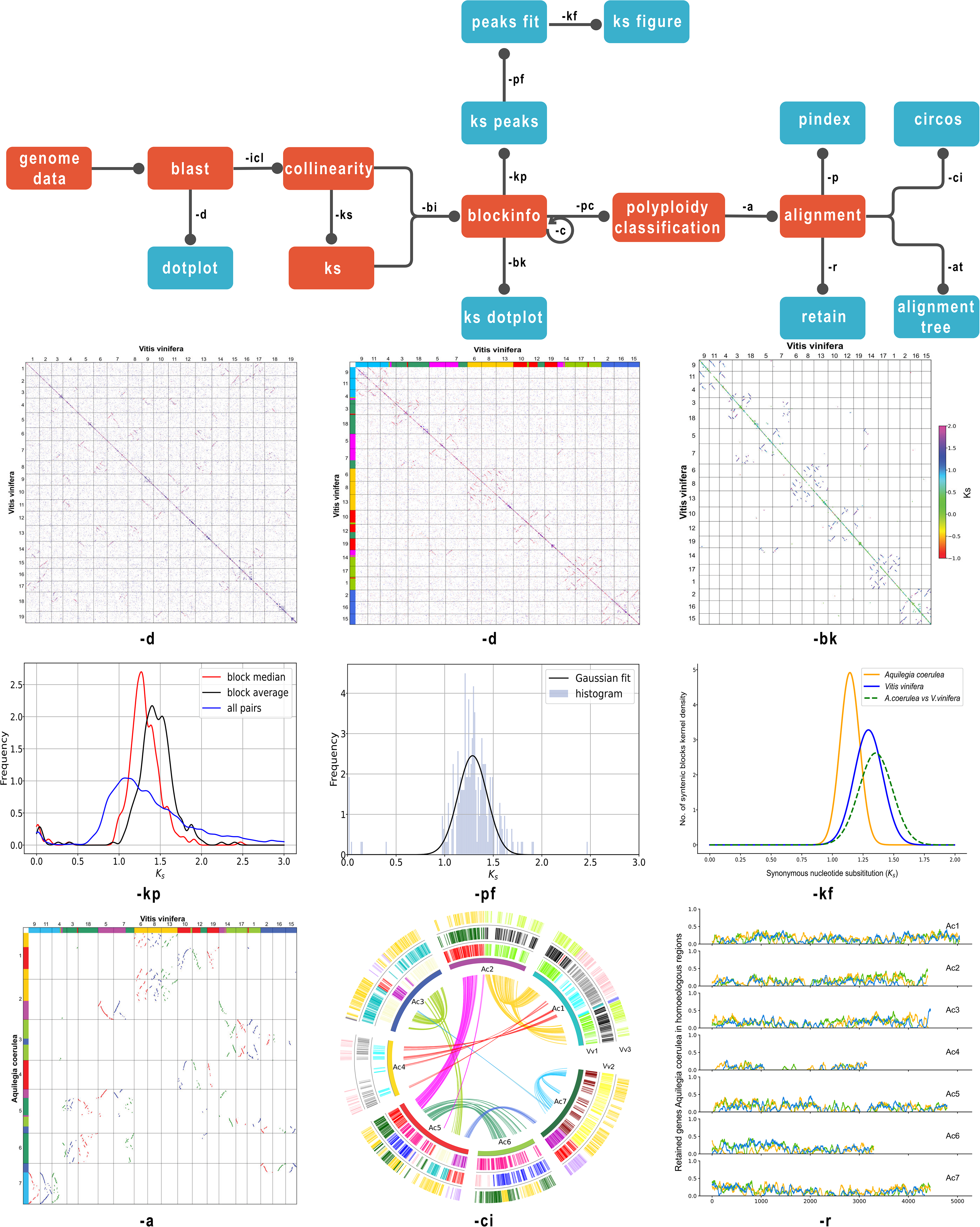
Structure of the WGDI package, illustrating major components and their dependencies.

### More sensitive collinearity detection

The synteny blocks extracted by WGDI are obviously accurate for two reasons. First, the settings of homologous genes are more flexible. WGDI only considers pairs of homologous genes in analyses, and retains homologous gene pairs related to polyploidization. This greatly reduces confounding effects of homology in large families (Supplementary Fig. 1). Second, WGDI scores and ranks homologous genes, following clear rules described in the homologous dotplot part of the Methods section, generating a dotplot with sets of red, blue, and gray dots with scores declining from high to low. For example, in block *a* − *c* shown in Fig. 2a, two dots are designated *b*_1_ and *b*_2_. If the homologous genes are not ranked and scored, the final scores will be the same, as illustrated by results obtained using the dynamic programming algorithm. In contrast, if they are ranked the gene with the higher homology (indicated here by the *b*_2_ dot) can be clearly identified.

**Fig. 2.**
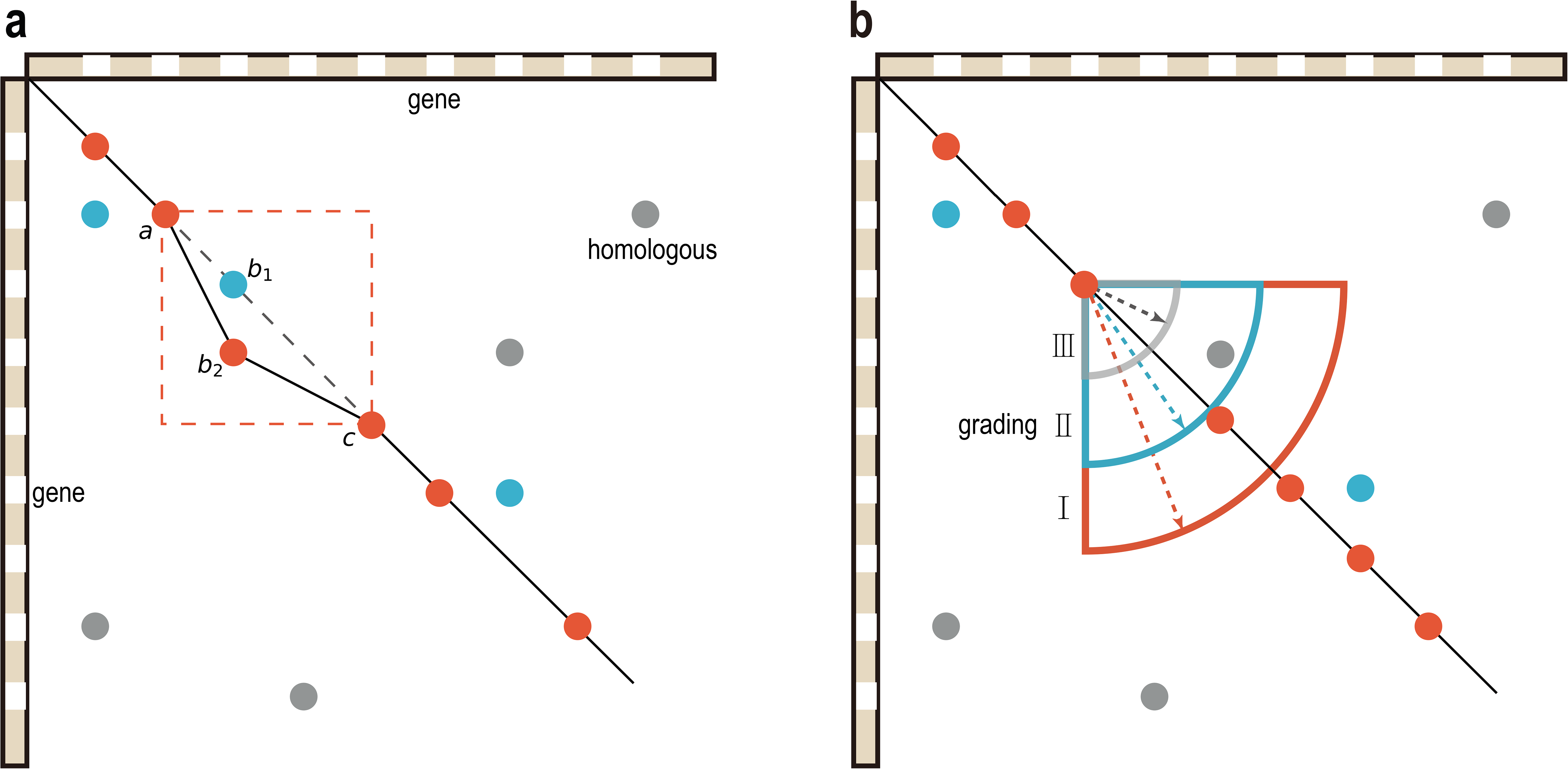
Flow chart of WGDI’s improved collinearity algorithm.

In addition, when the maximum gaps parameter is increased, the program will assign less highly homologous genes to the blocks, and the blocks will become longer. When the parameter is small, the error rate for ends of blocks will be reduced, but the blocks will be terminated earlier and shortened. After scoring and ranking homologous genes, the search range for homologous genes within blocks is also subject to a penalty rule, i.e. the range is positively related to the strength of homology (Fig. 2b). In this manner, the ends of blocks and maximum gap value can be optimized, and extracted blocks lengthened, without reducing sensitivity.

To evaluate the algorithm’s performance, we compared synteny blocks extracted by WGDI with those extracted by two other commonly used tools: MCScanX and JCVI (v1.1.12). The three tools were tested with the same datasets, Human/Chimpanzee (*Homo sapiens*/*Pan troglodytes*). The WGDI parameters were set to repeat_number=10, mg=25,25, muplite=1, grading=50,25,10, and score>100. MCScanX and JCVI parameters were set to default values. The number of synteny blocks extracted between *H. sapiens* chromosome 12 and *P. troglodytes* chromosome 13 by both WGDI and JCVI was 3, while that of MCScanX was 4. According to the dotplot on the left in Fig. 3, the explored region is rich in repeated genes. To illustrate capabilities of the three software packages for extracting synteny blocks in this region, we show a partial list of collinear genes in the figure, which we renamed according to their orders along the chromosome (the last five digits of each gene’s id indicate its position). Clearly, WGDI extracted more collinear genes than the other two packages, although JCVI performed better than MCScanX. In addition, both blast scores and gene arrangements suggested that synteny blocks extracted by JCVI and MCScanX included fewer or less accurate sets of homologous genes than those extracted by WGDI. We also compared synteny blocks in the region where the chromosome is inverted and drew similar conclusions (Supplementary Fig. 2).

**Fig. 3.**
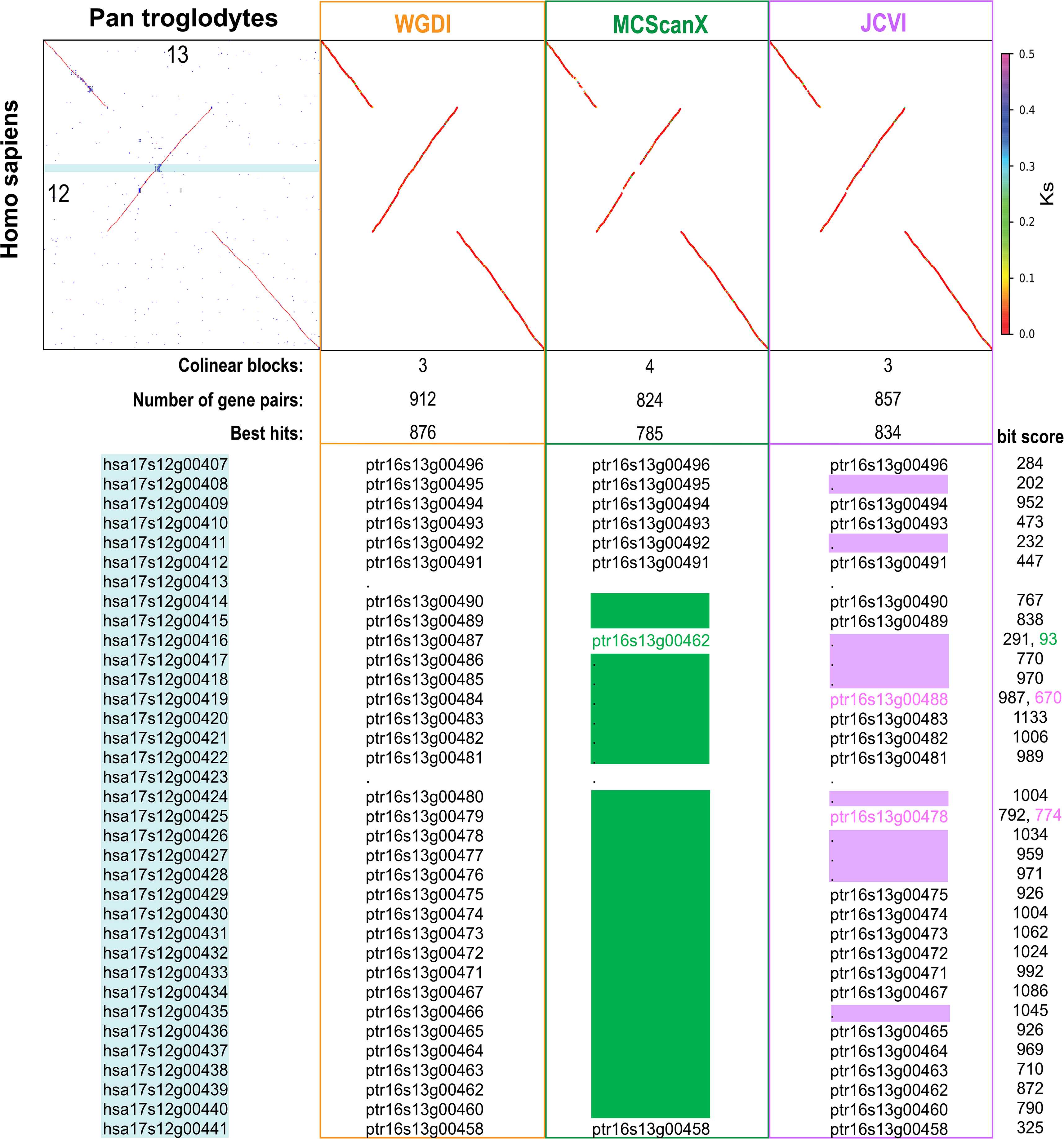
Comparison of the capability of WGDI, MCScanX, and JCVI for extracting synteny blocks.

### Examples of WGDI application

#### Polyploid inference

*Cucumis sativus* was selected as an example to illustrate the detailed process of polyploidy inference by WGDI. According to previous inferences, the *C. sativus* genome has undergone two rounds of polyploidization: the core-eudicot whole-genome triplication (WGT, or γ event) and a recent WGD (Huang, Li et al. 2009, Wang, Sun et al. 2018). *Vitis vinifera*, which has apparently only undergone one polyploidization (the γ event) in its evolutionary history, was selected as a reference species (Jaillon, Aury et al. 2007). The analysis involved the following three steps. First, the algorithm seeks evidence of WGDs by applying a synteny-based (homologous dotplot) method. In this case, the ‘−d’ program easily generated homologous *V. vinifera* versus *C. sativus* (Fig. 4a, Supplementary Fig. 3) and *C. sativus* vs *V. vinifera* (Fig. 4b, Supplementary Fig. 4) dotplots. In Fig. 4a, the longitudinal synteny depth is 3, which coincides with the γ event that occurred in all core dicots. The horizontal synteny depth may be 2, mainly because the red dots are not concentrated on one block. In Fig. 4b, the longitudinal synteny depth of the blocks mainly consisting of red dots is 2. In combination with three homologous chromosomes of *V. vinifera*, the maximum synteny depth can reach 3*2 = 6. The horizontal synteny depth of the blocks mainly comprising red dots is 1. The maximum synteny depth is 3*1=3. Combining the known WGDs and synteny depth, the evolutionary tree diagram can be inferred (Fig. 4c). Therefore, the homologous dotplot can easily implement most WGD speculations.

**Fig. 4.**
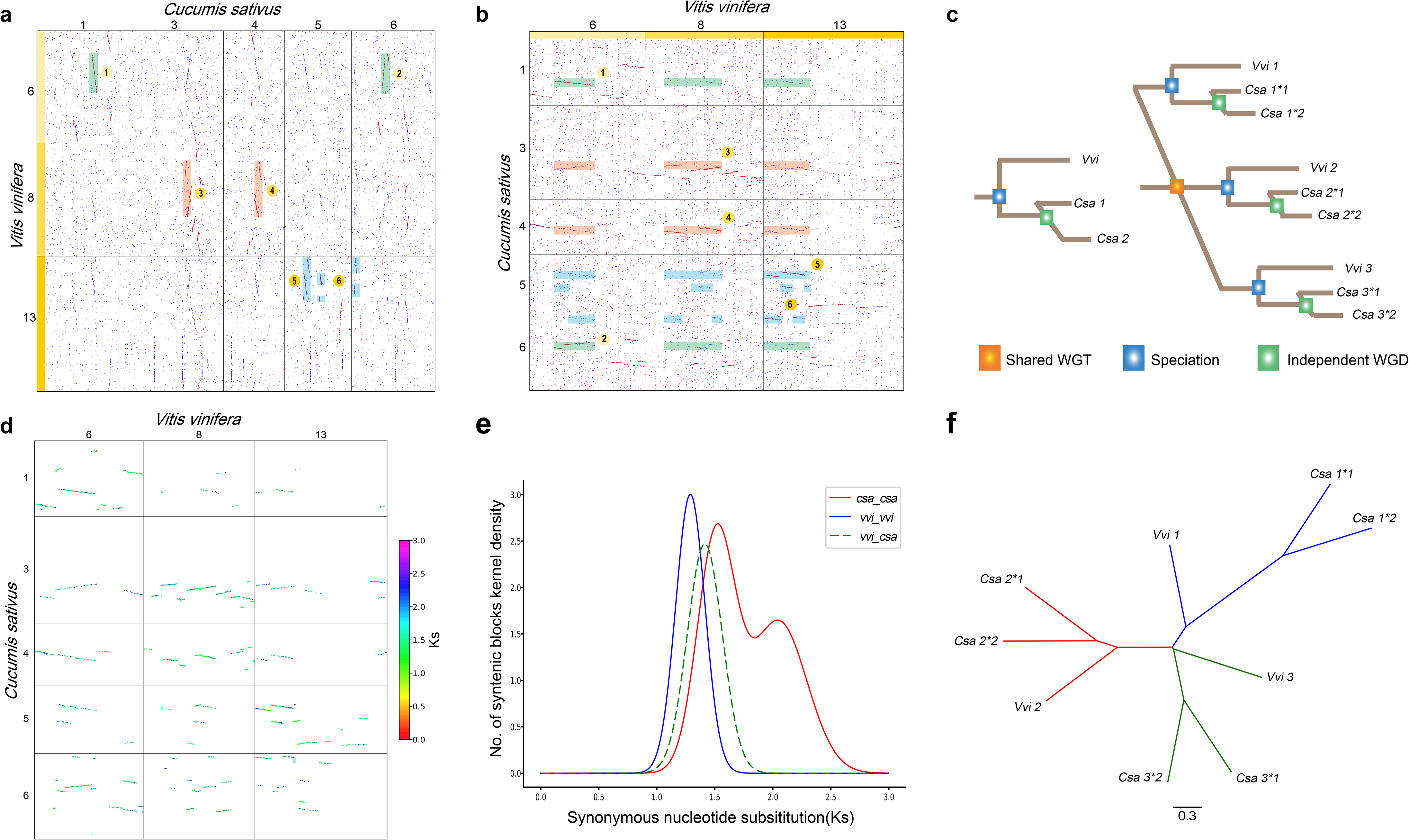
Polyploid inference using *Cucumis sativus* and *Vitis vinifera* as examples. (a) Homologous dotplot between selected *V. vinifera* and *C. sativus* chromosomes. The *V. vinifera* chromosomes 6, 8, and 13, being homoeologous triplets produced by the core-eudicot whole-genome triplication. (b) Homologous dotplot between selected *C. sativus* and *V. vinifera* chromosomes. Outparalogous regions or secondary-matched were marked out by rectangles. (c) Gene phylogenetic trees of these two species. (d) Ks dotplot. Syntenic blocks were identified and colored based on their Ks values. (e) Ks distribution. (f) Phylogenetic relationships inferred by the alignment trees using ASTRAL-III.

Secondly, the Ks distribution is used to verify the occurrence and times of WGDs. Through the ‘−bk’ program, the Ks value of gene pairs on synteny blocks can be displayed intuitively (Fig. 4d). The Ks value of gene pairs on the same block fluctuates slightly, and the median can be used to represent the Ks distribution of the block. The Ks value of different blocks fluctuates greatly, especially when the blocks were retained after different polyploid events. Therefore, combining the *Homo* parameter and the Ks distribution, it is easy to distinguish these blocks, and then output the fitting function and the best fit through the ‘−pf’ program. Currently, many researchers are still using Gaussian mixture models or similar methods to fit multiple polyploid events. Since the Gaussian mixture model is prone to overfitting, the Ks peak obtained is fluctuating and inaccurate (Zwaenepoel and Van de Peer 2018). The Ks distribution within or among multiple species can be displayed through the ‘−kf’ program. It can be seen that the Ks peaks of the triploidization event shared by *V. vinifera* and cucumbers are very different, and even the Ks peaks of the differentiation events of *V. vinifera* and cucumbers are smaller than the Ks peaks of the recent WGD of cucumbers (Fig. 4e). Due to the great differences in evolutionary rates of species, the Ks peaks of the same WGD may be quite different. Therefore, it is essential to correct Ks peaks if necessary, and then infer the time of polyploidy (Wang, Sun et al. 2017, Wang, Sun et al. 2018, Yang, Sun et al. 2020). Compared with the method of paralogous gene pairs obtained through clustering methods (such as OrthoMCL), the Ks distribution inferred by WGDI could largely reduce the effect of tandem duplication or other non-syntenic paralogous.

Finally, the gene tree-based method is used to effectively verify polyploidy inference. Through the “-a” and “at” programs, the homology list of *V. vinifera* and *C. sativus* was obtained, the phylogenetic relationship was inferred from 1138 of the constructed gene trees by ASTRAL-III v5.7.7 (Zhang, Rabiee et al. 2018) (Fig. 4f). Among them, the phylogenetic relationship was inferred from 92 trees that have at least 3 cucumber genes is the same, accounting for 8.08%. Due to gene loss, the structure of trees that have lost several homologous genes supports that *Cucumis* experienced a more recent WGD. In short, WGDI brings together three main methods to detect WGD and verifies each other to help researchers complete polyploid inference accurately and efficiently.

#### Hierarchical inference of genomic homology

*Arachis hypogaea* is an allopolyploid that has been shown to originate from a cross between two ancestral species, *A. duranensis* and *A. ipaensis* (Bertioli, Cannon et al. 2016, Bertioli, Jenkins et al. 2019, Zhuang, Chen et al. 2019). Here we use *A. hypogaea* as an example to show how to infer genomic homology of the allopolyploid by the following steps. Firstly, homologous dotplots and Ks dotplots are used for hierarchical reasoning of genome homology. The ‘−d’ and ‘−bk’ programs suggest that chromosomes 1-10 of *A. hypogaea* are derived from *A. duranensis* and chromosomes 11-20 from *A. ipaensis*. However, there has been clear translocation between chromosomes 3 and 13 (Fig. 5a). According to differences in synteny blocks retained by different polyploid events, including Ks, the identity of blocks, *Homo* and others, it is easy to obtain the list of homologous genes among the three genomes through the ‘−a’ program (Fig. 5b). Secondly, the alignment tree is used for further hierarchical inferences. Through the ‘- at’ program, the homology list of *A. hypogaea* chromosome 3 with translocation and those of *A. duranensis* and *A. ipaensis* is used to construct 2,376 gene trees, with all possible tree structures scored and displayed in the form of sliding windows (Fig. 5c). Among them, 1597 trees support origination of *A. hypogaea* from *A. duranensis* and 624 support its origination from *A. ipaensis*, which is consistent with the inferred chromosomal translocation.

**Fig. 5.**
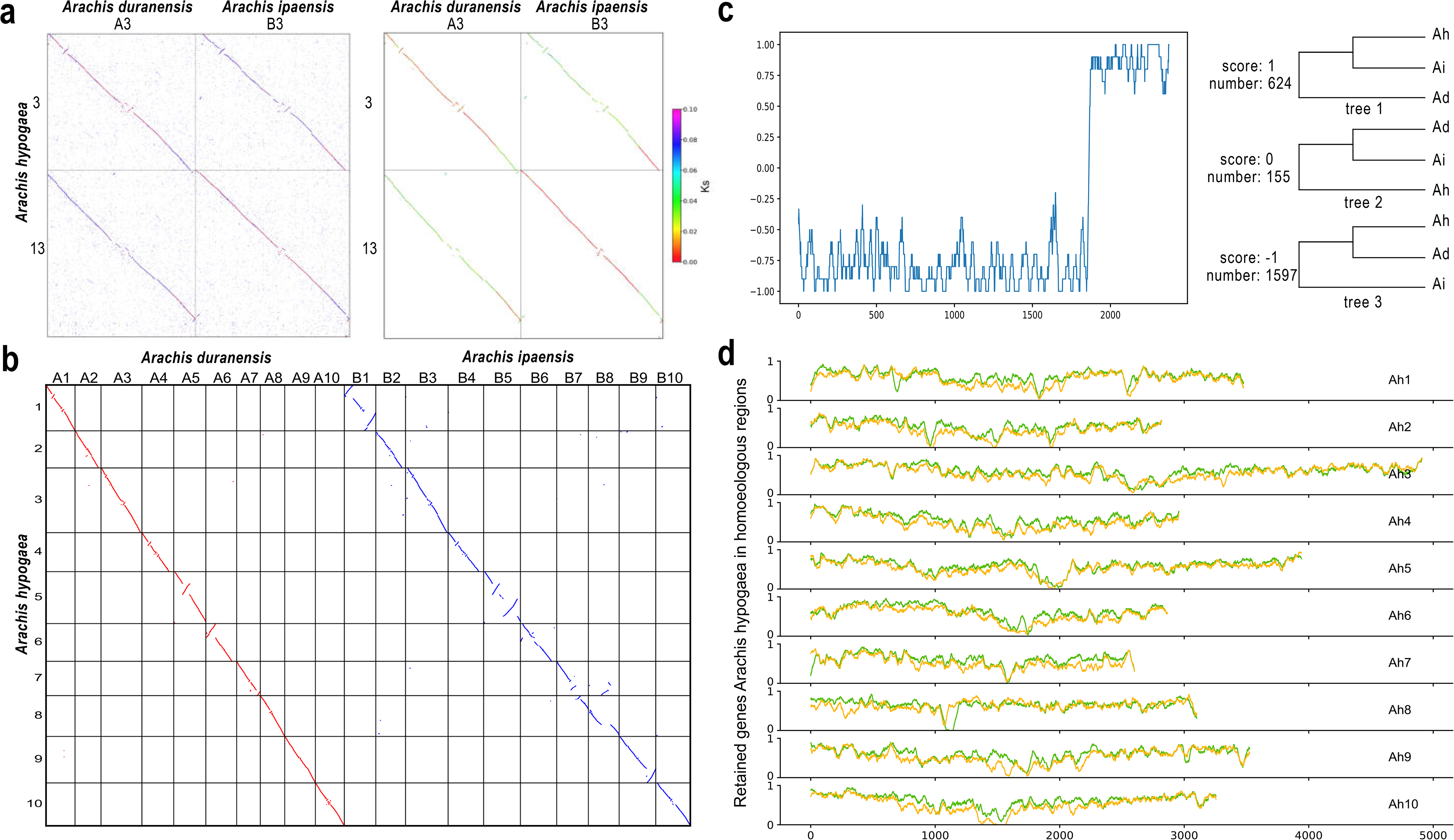
Hierarchical inference of genomic homology using *Arachis hypogaea* as examples. (a) Homologous dotplot and Ks dotplot between selected *A. hypogaea* and *A. duranensis* and *A. ipaensis* chromosomes. The *A. hypogaea* chromosomes 3 and 13, being homoeologous triplets produced by the eudicot-common hexaploidy, and their matched *A. duranensis* chromosomes 3 and *A. ipaensis* chromosomes 4. (b) Alignment of *A. hypogaea* with *A. duranensis* as a reference genome. (c) Distribution of all possible tree structures in the homology list of *A. hypogaea* chromosome 3. (d) Rates of retained genes in sliding windows of ah homologous region group 1 (orange) and homologous region group 2 (green).

Finally, through the ‘−r’ and ‘−p’ programs, the gene retention and subgenome divergence in the allopolyploid are respectively displayed. In the allotetraploid *A. hypogaea*, more genes from the *A. duranensis* subgenome were more retention than those from the *A. ipaensis* subgenome (Fig. 5d). In addition, the divergence (P-index) between two subgenomes or two ancestral diploid genomes was estimated as 0.86, indicating substantial difference between them, as in a typical allotetraploidization. Therefore, WGDI uses multiple methods to perform hierarchical inferences of allopolyploidization based on genome homology and generates a hierarchical list of homologous genes supported by gene collinearity and related to allopolyploid speciation.

#### Ancestral chromosomal karyotype

The chromosomal karyotype of *V. vinifera* is widely used as a reference for karyotype analyses of core eudicots. The previous karyotype analysis suggested that the ancestral chromosomes of *V. vinifera* experienced two chromosomal fusions and four chromosomal fissions (Murat, Zhang et al. 2015). Based on karyotype changes, it was speculated that *Aquilegia coerulea* (Ranunculaceae) was one parental ancestor of all core dicots as a likely ancient allohexaploid (Aköz and Nordborg 2019). However, through the homologous dotplot of *V. vinifera* and *A. coerulea* and the ancestral karyotype of *A. coerulea*, a lot of short synteny blocks appear bizarrely in a region that does not belong to it (Fig. 6a, Supplementary Fig. 5). These blocks are verified by synteny analysis and Ks analyses (Supplementary Fig. 6). The assembly of multiple versions of *V. vinifera* genomes indicates that these fragments do exist (Supplementary Fig. 7-8). For example, the ends of *A. coerulea* Chr2 and Chr3 and the front of Chr5 correspond to the middle region of *V. vinifera* Chr4 (Fig. 6a), which is an unlabeled region (blank area, see Fig. 6 in Aköz and Nordborg 2019). Such results may be misleading if karyotype evolution derived from these blocks is not considered.

**Fig. 6.**
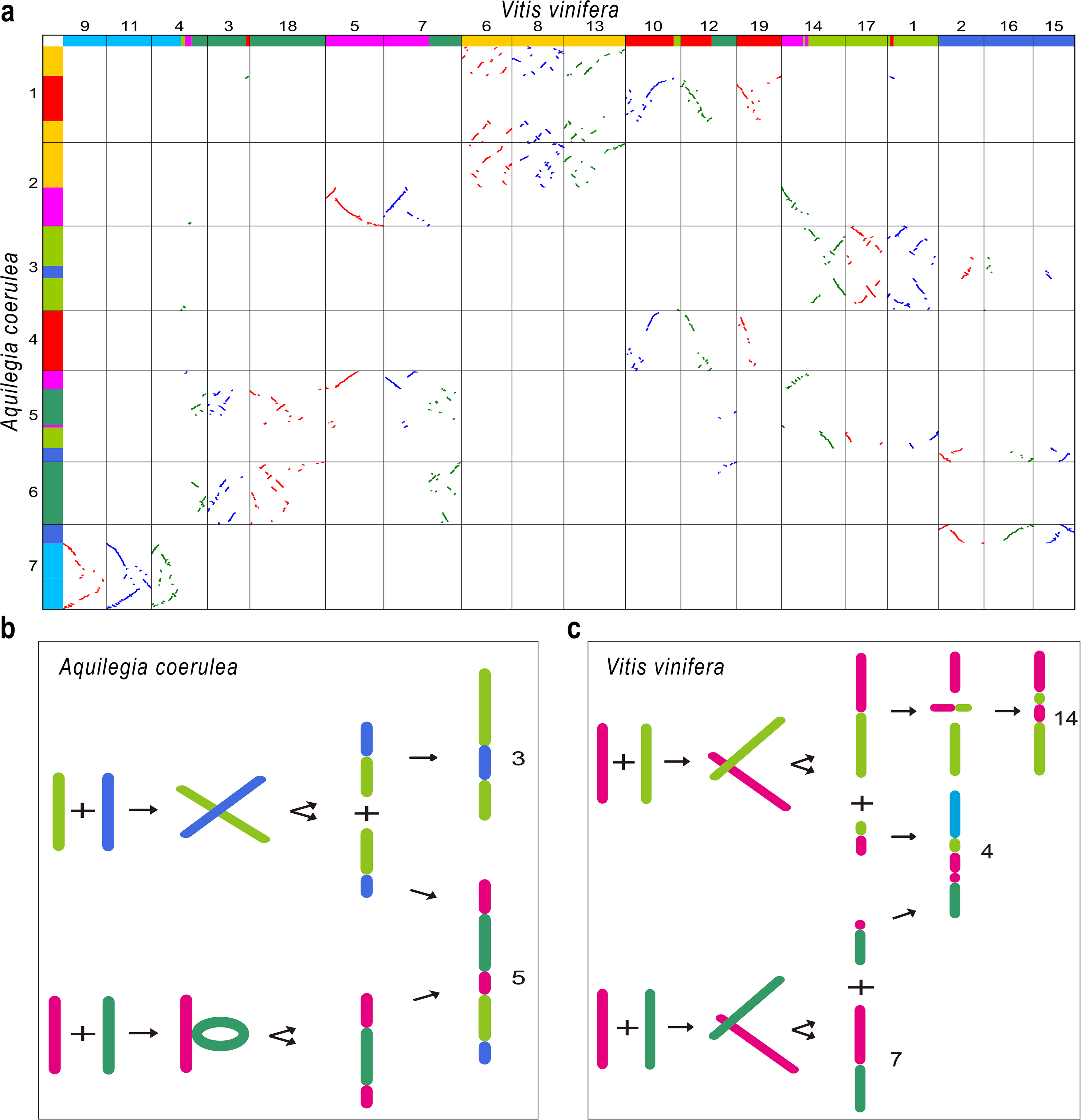
Karyotype evolution of *Vitis vinifera* chromosomes 4, 7 and 14 and *Aquilegia coerulea* chromosomes 3 and 5. (a) Homologous dotplot between selected A. coerulea and V. vinifera. Seven distinct colors represent the haploid set of seven ancestral chromosomes. (b) Process of construction of A. coerulea chromosome 3 and 5. (c) Process of construction of V. vinifera chromosome 4, 7 and 14.

The chromosome number reduction and B chromosome model (Wang, Jin et al. 2015, Wang and Wang 2020) provide new methods for understanding the evolution of eukaryotic chromosome numbers and chromosome rearrangement. According to this model, the ancestral karyotype of *V. vinifera* is more complex but clearer than previously inferred. For example, pink and grass-green chromosomes crossed over near one of two ends of each chromosome to produce two novel chromosomes (Fig. 6c). Among them, two long parts fused into the *V. vinifera* Chr14 and inverted later within this chromosome. The two short parts did not disappear, but fused with other chromosomes to form the *V. vinifera* Chr4. Pink and green chromosomes underwent a similar fusion, leading to the current Chr7 and Chr4. However, the karyotype of *A. coerulea* is completely different from that of *V. vinifera*. The green chromosome was inserted into the pink chromosome and the grass-green chromosome and blue chromosome crossed over, then evolved into the current Chr5 and Chr3 (Fig. 6b). Therefore, karyotypes of *A. coerulea* and *V. vinifera* (Aköz and Nordborg 2020) cannot be used to support the conclusion that the *Aquilegia* genome reveals a hybrid origin of core eudicots. Compared with previous conclusions, our new conclusion revealed by WGDI may better reflect the dynamic process of ancestral chromosomal rearrangement. Such blocks also exist in other species, and most of them are ignored without detailed discussion.

## Discussion

Polyploidization is recognized as an important driving force for the evolution of species. It plays an important role in the evolution of species and formation of new species. Gene collinearity is an important way to study species polyploidization. Although several tools for analyzing multiplication have emerged in recent years, there have been few substantial improvements in the algorithm for collinearity extraction, and the downstream evolution analysis program provided no distinction between collinearity fragments caused by multiplication. This incompleteness of functionality has reduced the usefulness of existing collinearity detection tools. WGDI is particularly useful for identifying polyploidy events, and the inference of hierarchy and event-related gene collinearity proposed by this tool helps the actual phylogeny of plants affected by recursive polyploidization. Also, many biological analyses implemented in WGDI are unique. WGDI outperforms similar tools in terms of efficiency, flexibility, scalability.

In addition, WGDI is highly useful and effective for reconstructing ancestral chromosomal karyotype of current species. WGD causes rapid genome reorganization and structural variations to produce the new chromosomal karyotype of the following lineage or species. Such karyotype evolution is regarded as an important factor for evaluating the phylogenetic position of one disputed lineage and inferring the genome structure of an extinct species (Aköz and Nordborg 2020). However, two models of fission and fusion obviously (Salse, Bolot et al. 2008, Murat, Xu et al. 2010) cannot understand how chromosomes of the ancestral genome evolved into the current karyotypes (Wang, Jin et al. 2015). For example, the ancestral karyotypes of all angiosperms, monocots, and core dicots (Murat, Armero et al. 2017) have numerous large unlabeled (blank) regions. The shared synteny or synteny breaks during karyotype evolution are critical characters for such inference and a more rigorous framework to perform such analyses is badly needed in addition to the simple fission-fusion model (Aköz and Nordborg 2020). WGDI can accurately extract syntenic blocks and thus facilitate reconstruction of the evolutionary process of ancestral chromosomes.

## Conclusions

WGDI outperforms similar tools in terms of efficiency, flexibility and scalability in WGD evolutionary analyses. This new toolkit implements a dynamic programming-based collinearity extraction algorithm and incorporates multiple computer programs for visualization and analysis. Inference of hierarchical and event-related gene colinearity helps actual phylogeny of affected by recursive polyploidization. WGDI is freely available for public use via GitHub (https://github.com/SunPengChuan/wgdi). WGDI also makes use of conda environments and the bioconda platform (Grüning, Dale et al. 2018), which allows hassle-free installation and upgrading of known-compatible and known-functional tools.

## Methods

### Homologous dotplot

The protein-coding genes from each genome were compared against itself and other genomes using BLASTP (E-value < 10-5, and Score > 100) or other three softwares, BLAST+ (Camacho, Coulouris et al. 2009), MMseqs2 (Steinegger and Söding 2017) and DIAMOND (Buchfink, Reuter et al. 2021). According to the descending score order, the number of unique homologous genes is restricted for a sequence. The parameter ‘multiple’ is set to the best number of homologous genes, and these gene pairs are shown with red dots. The next four homologous gene pairs are shown in blue dots with the rest in gray dots. ‘multiple’ is closely related to polyploidy among species.

### Improved collinearity

The algorithm for extracting synteny blocks is based on a dynamic programming algorithm, similar to ColinearScan (Wang, Shi et al. 2006) and MCScan (Tang, Wang et al. 2008). Assuming that two gene pairs, *u* and *v*, are on the path where *u* precedes *v*,

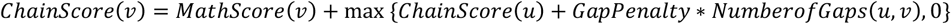

*MathScore(v)* assigns different scores based on the colors in the dotplot, with a default of 50 for red, 40 for blue, and 25 for gray. These parameters can be easily adjusted. *GapPenalty* = −1, and *NumberofGaps*(*u*, *v*) the maximum number of intervening genes between *u* and *v*, defaults to 40. These parameters can be modified directly in the configuration file. The minimum collinear block is 5 collinear gene pairs. To evaluate the compactness and uniqueness of collinear blocks, we set up the ‘pvalue’ parameter.

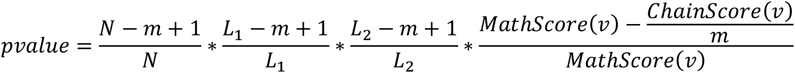

Here, N is the number of genes in the coverage [*v* − 0.5 ∗ *NumberofGaps*(*v*), *v* + 0.5 ∗ *NumberofGaps*(*v*)] of any genes on the collinear path. *L*_1_, *L*_2_ are the respective lengths of the two chromosomal regions, and *m* is the number of colinearity pairs. In order to calculate the ratio of the best homology on the collinearity block, we set the parameter *Homo*, that is, the red dots in the homologous dotplot are marked as 1, the blue dots are marked as 0, and the gray dots are marked as −1. Then,

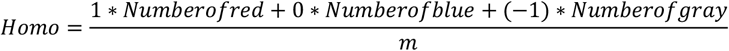

In this way, the *Homo* range of the red block in the dotplot is roughly [0.5, 1, and the blue block is [-0.5, 0.5. By integrating *Homo*, collinearity results and Ks results into a ‘blcokinfo’ file, we can combine multiple screening conditions to screen collinearity fragments generated by different polyploid events.

### Non-synonymous (Ka) and synonymous (Ks)

Protein sequences of colinear homologs inferred above were aligned using MUSCLE (Edgar 2004) or MAFFT (Katoh, Rozewicki et al. 2019). The resulting protein alignments are used to guide coding sequence alignments by PAL2NAL (v14) (Suyama, Torrents et al. 2006). Non-synonymous (Ka) and synonymous (Ks) substitution rates are estimated using the Nei-Gojobori method implemented in the YN00 program in the PAML (4.9h)(Yang 2007) package. Note that some homologous gene pairs are assigned with a value of −1 due to misalignment.

### Inference of hierarchical and event-related gene collinearity

In order to clarify the multi-level collinearity fragments caused by recursive polyploidization, WGDI combines the dotplot, collinearity, and Ks values to generate a table file that records the index information of each collinearity block. In this way, it is quick to filter and statistically plot certain indicators. WGDI can classify each collinearity segment to infer hierarchical and event-related gene collinearity.

### Interactive visualization

WGDI provides a visual way to place the Ks value in the synteny block on the dotplot. After screening the collinearity fragments, the Ks distributions (median, average, total) in three ways are displayed through nuclear density analysis. WGDI can group these collinearity fragments according to polyploid events or sub-genomes to generate a list related to polyploid events and visualize the extracted collinearity. WGDI also provides a circos drawing form to show this collinearity list.

### Retain and alignment tree

WGDI visualizes alignment, that is it displays the level of gene retention between reference genome chromosomes, subgenomes, or different species. Because the alignment genes are all extracted from collinearity genes and related to polyploid events, using these gene sets to call IQTREE(Minh, Schmidt et al. 2020) to construct a phylogenetic tree to show the actual phylogenetic relationship.

### Polyploidy index

The polyploidy index (P-index) (Wang, Qin et al. 2019) is used to characterize the degree of divergences among subgenomes of a polyploid species and find whether balanced or unbalanced genes were removed from homologous regions. The parameter “retention” is designed to remove the region where the retention rate of the sub-genome relative to the reference genome is low.

### Availability

WGDI is freely available for public use at GitHub (https://github.com/SunPengChuan/wgdi). Documentation for installation along with a user tutorial (https://wgdi.readthedocs.io/en/latest/index.html), default parameter file, and test data at GitHub(https://github.com/SunPengChuan/wgdi-example) are provided. Additional datasets analyzed in the current study are available upon request from the corresponding author.

## Acknowledgments

We thank Drs. Chengjie Chen and Zhougeng Xu for their contributions in software promotion.

## Author contributions

J.L., X.W. and Z.X. supervised and managed the project and research. P.S. and B.J. finished most analyses and tests. Y.Y., L.S., T.L., and X.L. participated in software testing and provided suggestions for revisions. P.S. and B.J. wrote the manuscript. J.L revised the manuscript and X.W. and Z.X. are involved in the discussion and improvement of the manuscript. All authors read and approved the final manuscript.

