## Supplemental for "WGDI: A user-friendly toolkit for evolutionary analyses of whole-genome duplications and ancestral karyotypes"

Supplementary Figures ........................................ 2

### Supplementary Figures

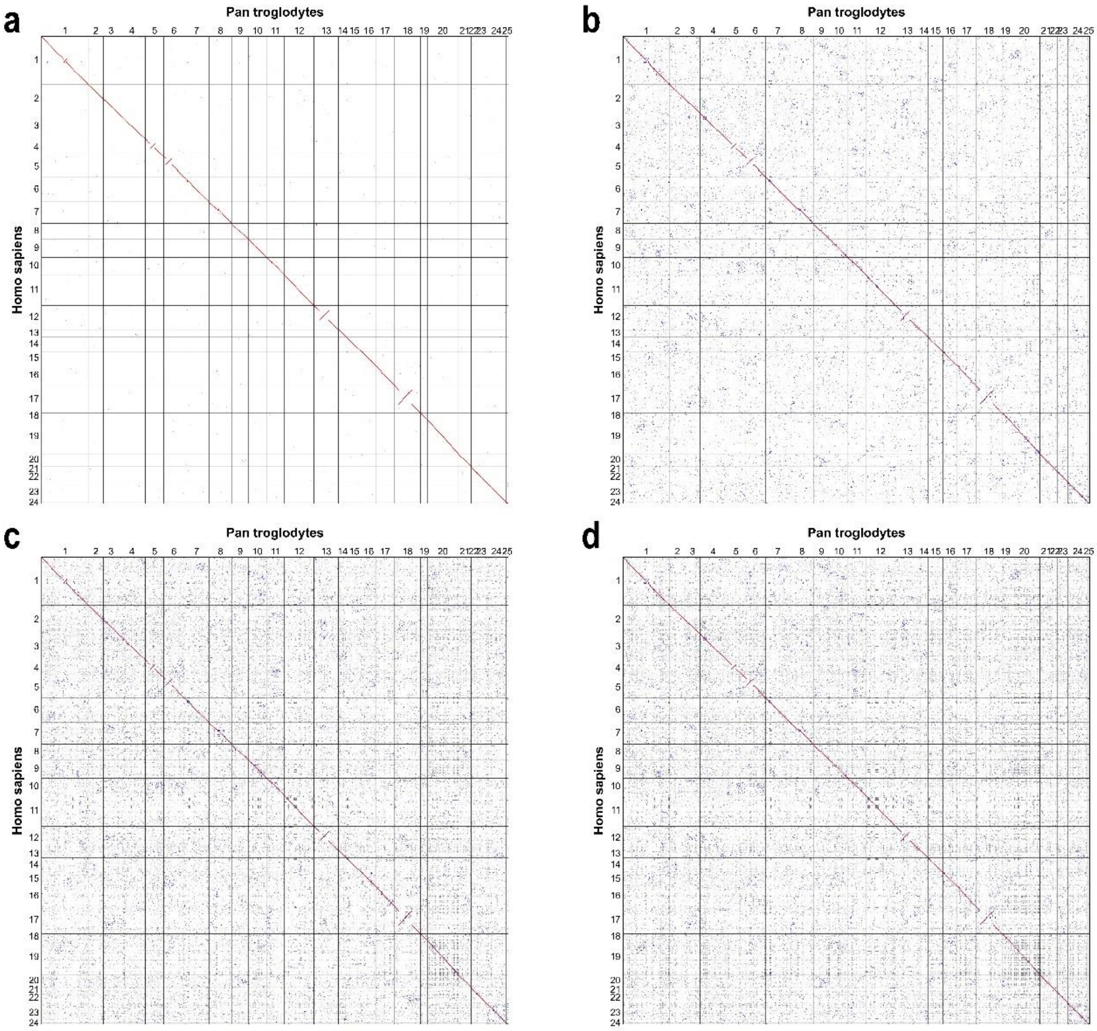


**Supplementary Fig. 1 Homologous dotplot between *Homo sapiens* and *Pan troglodytes.*** (a) repeat_number = 1. (b) repeat_number = 10. (c) repeat_number = 100. (d) repeat_number = Max.

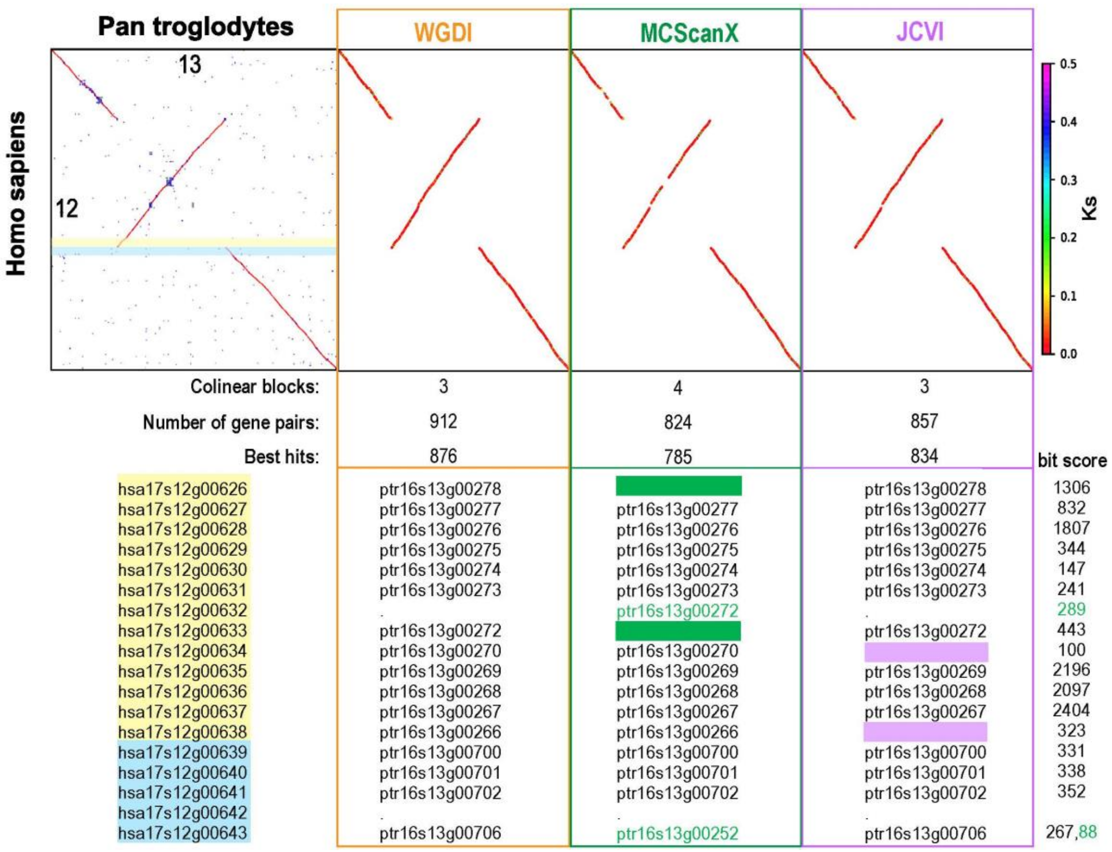


**Supplementary Fig. 2 Comparison of the capability of WGDI, MCScanX, and JCVI for extracting synteny blocks.**

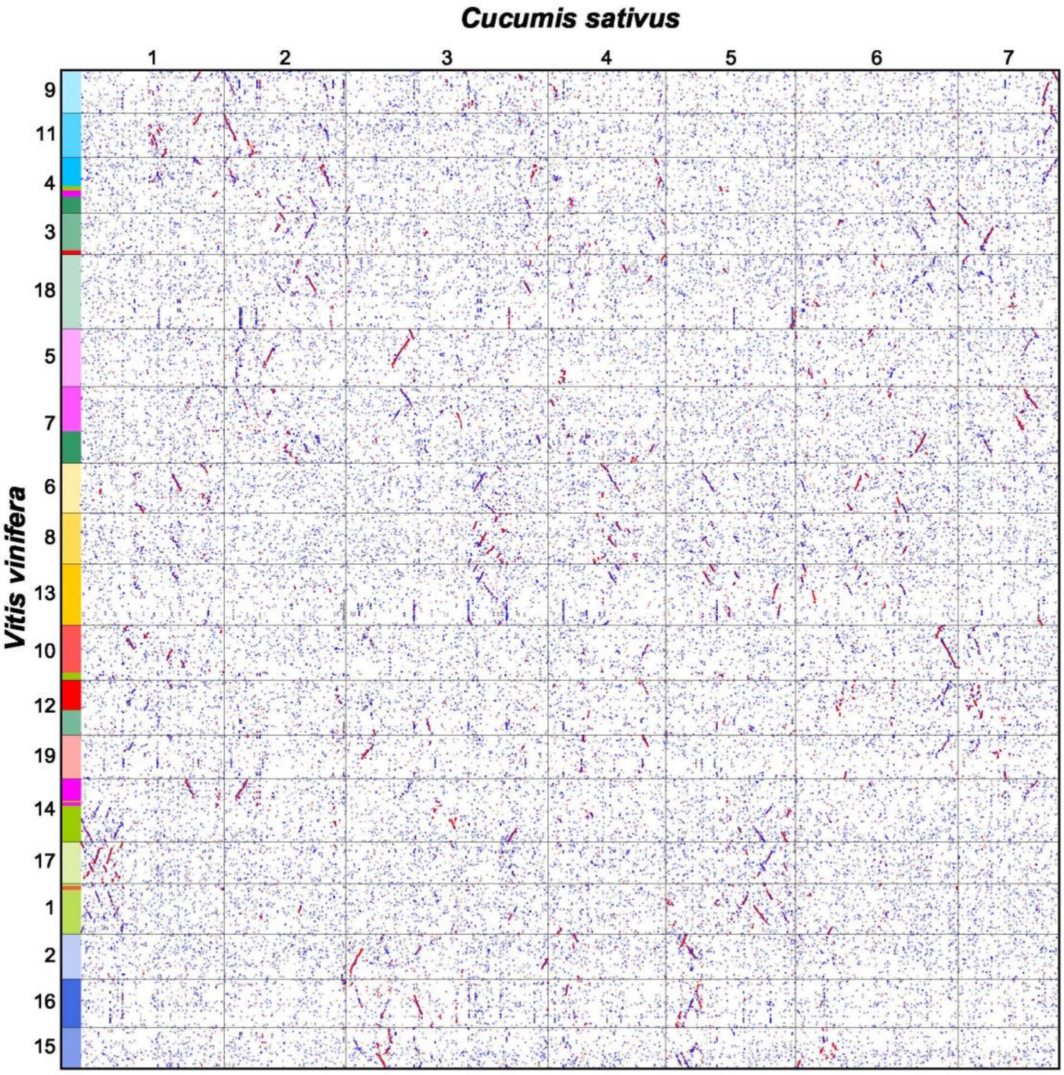


**Supplementary Fig. 3 Homologous dotplot between *Vitis vinifera* and *Cucumis sativus*. The color-coding of the 19 chromosomes of grapes corresponds to the seven ancestral chromosomes of core dicots. The parameters are Multiple = 1 and repeat_number = 10.**

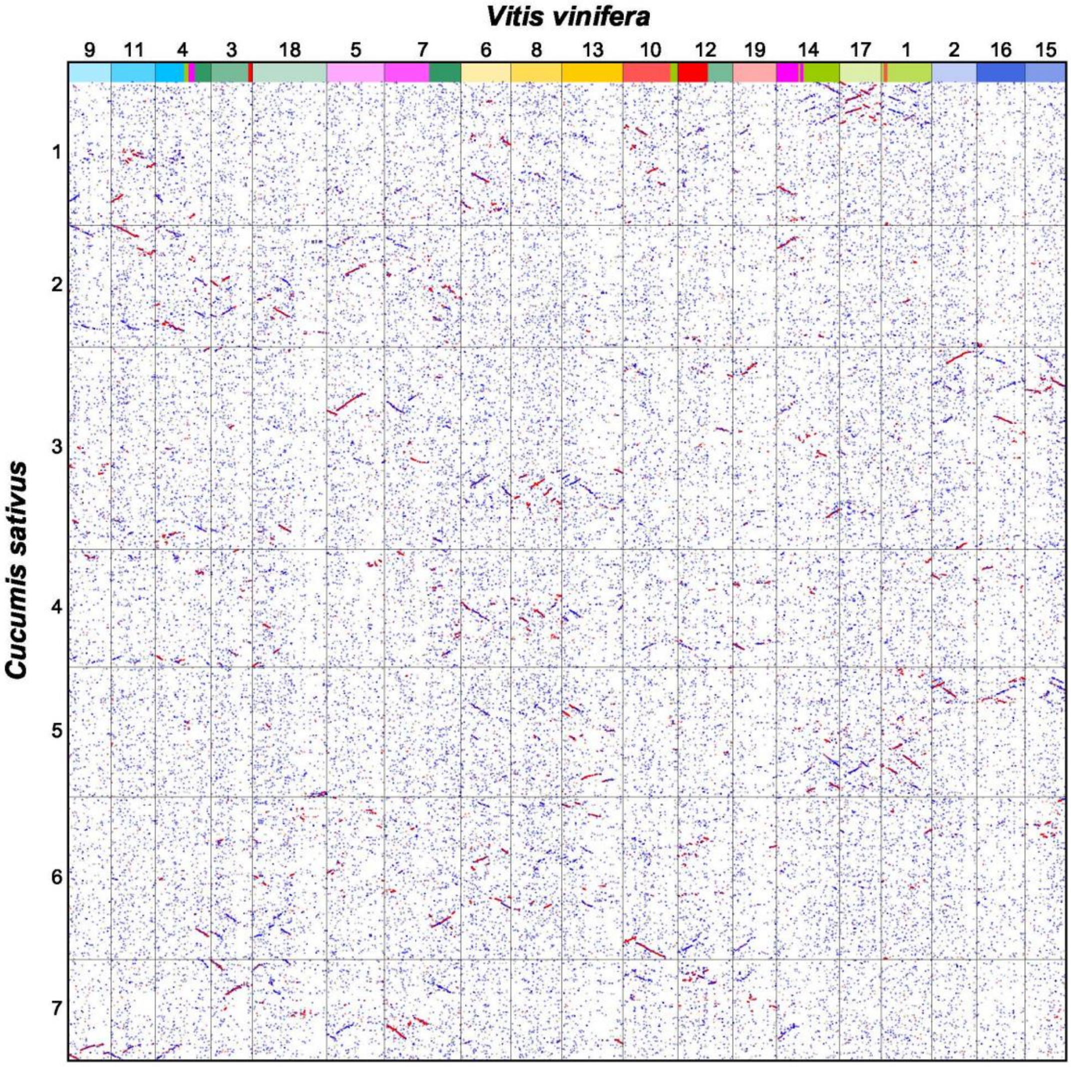


**Supplementary Fig. 4 Homologous dotplot between *Cucumis sativus* and *Vitis vinifera.*** The color-coding of the 19 chromosomes of grapes corresponds to the seven ancestral chromosomes of core dicots. The parameters are Multiple = 1 and repeat_number = 10.

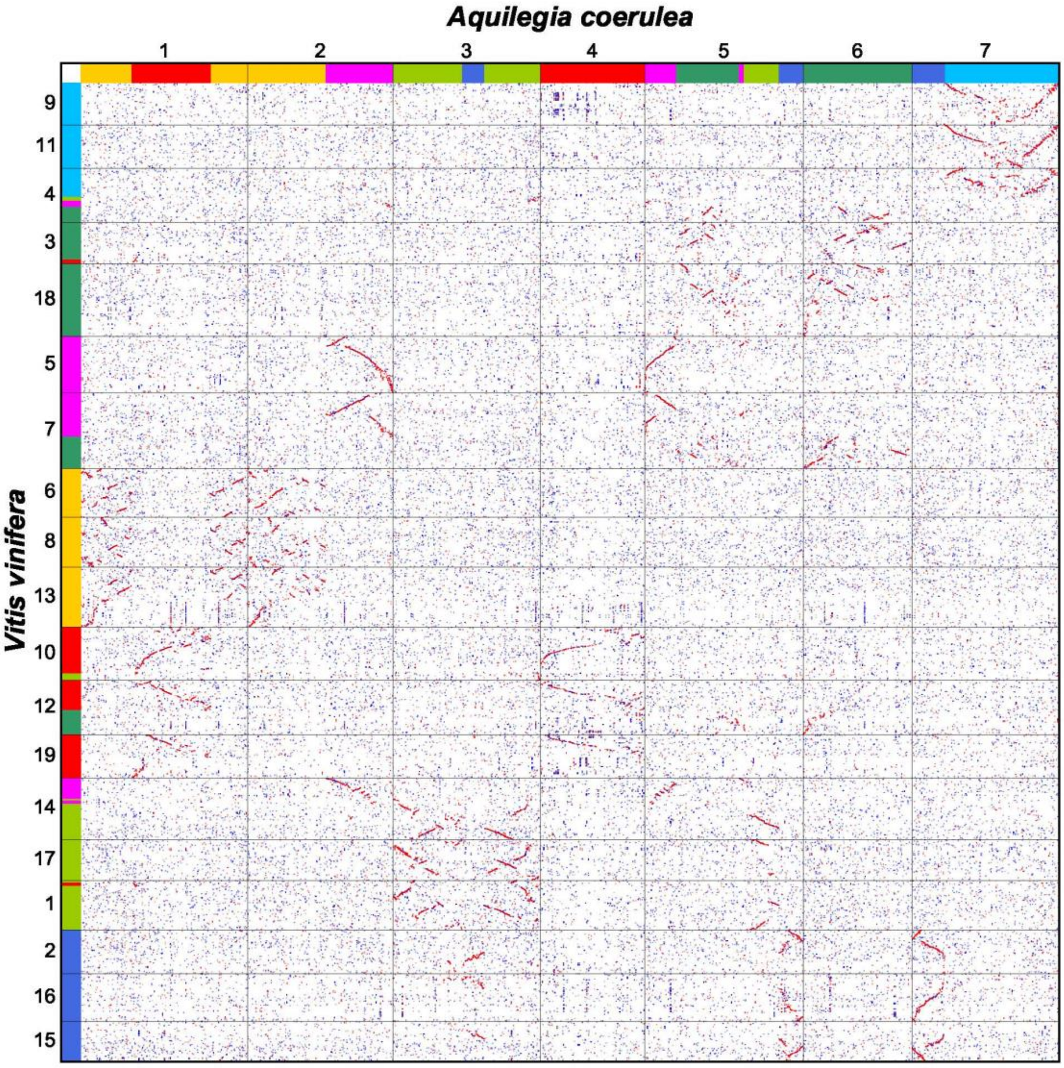


**Supplementary Fig. 5 Homologous dotplot between *Vitis vinifera* and *Aquilegia coerulea.*** The color-coding of the chromosomes of *V. vinifera* and *A. coerulea* corresponds to the seven ancestral chromosomes of core dicots. The parameters are Multiple = 1 and repeat_number = 10.

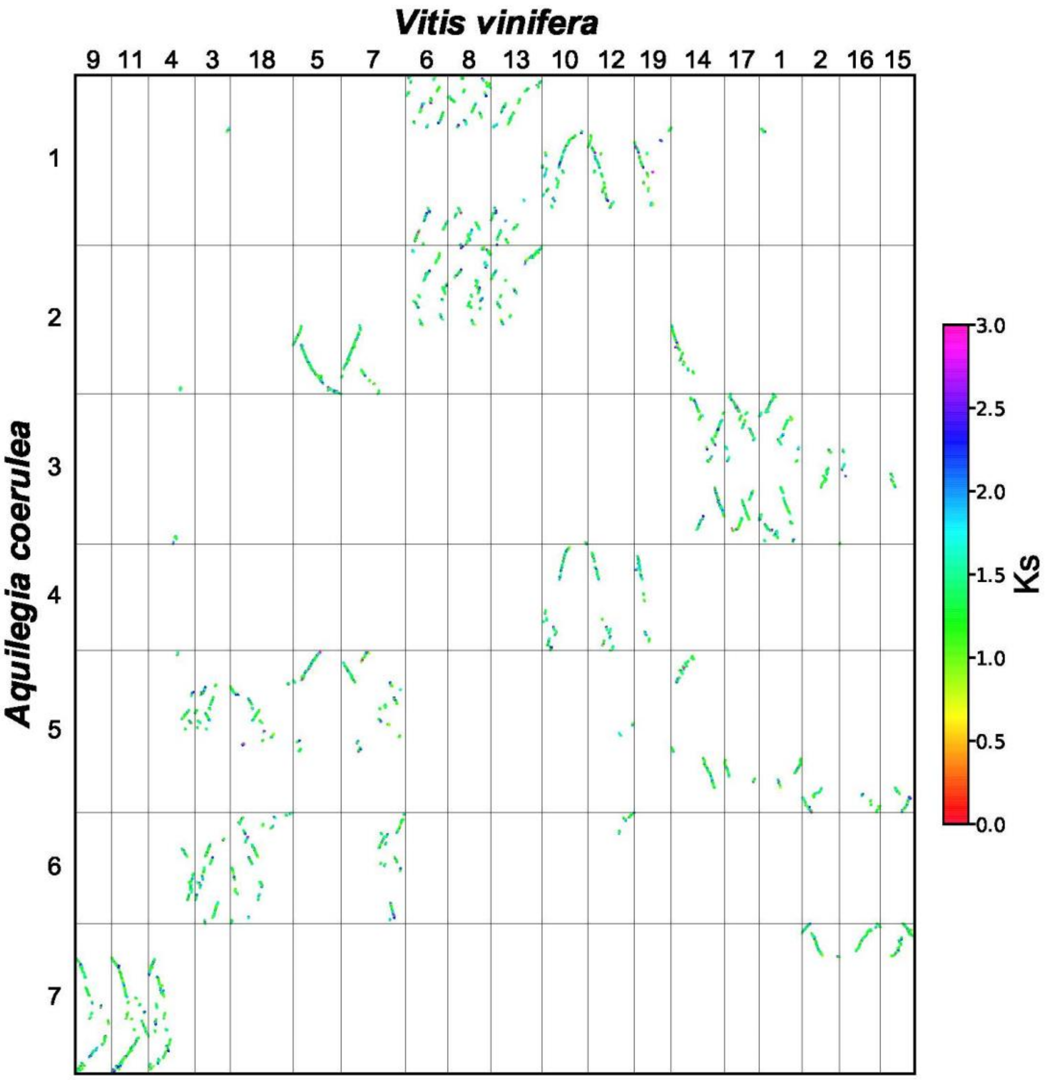


**Supplementary Fig. 6 Syntenic blocks within *Vitis vinifera* and *Aquilegia coerulea.***

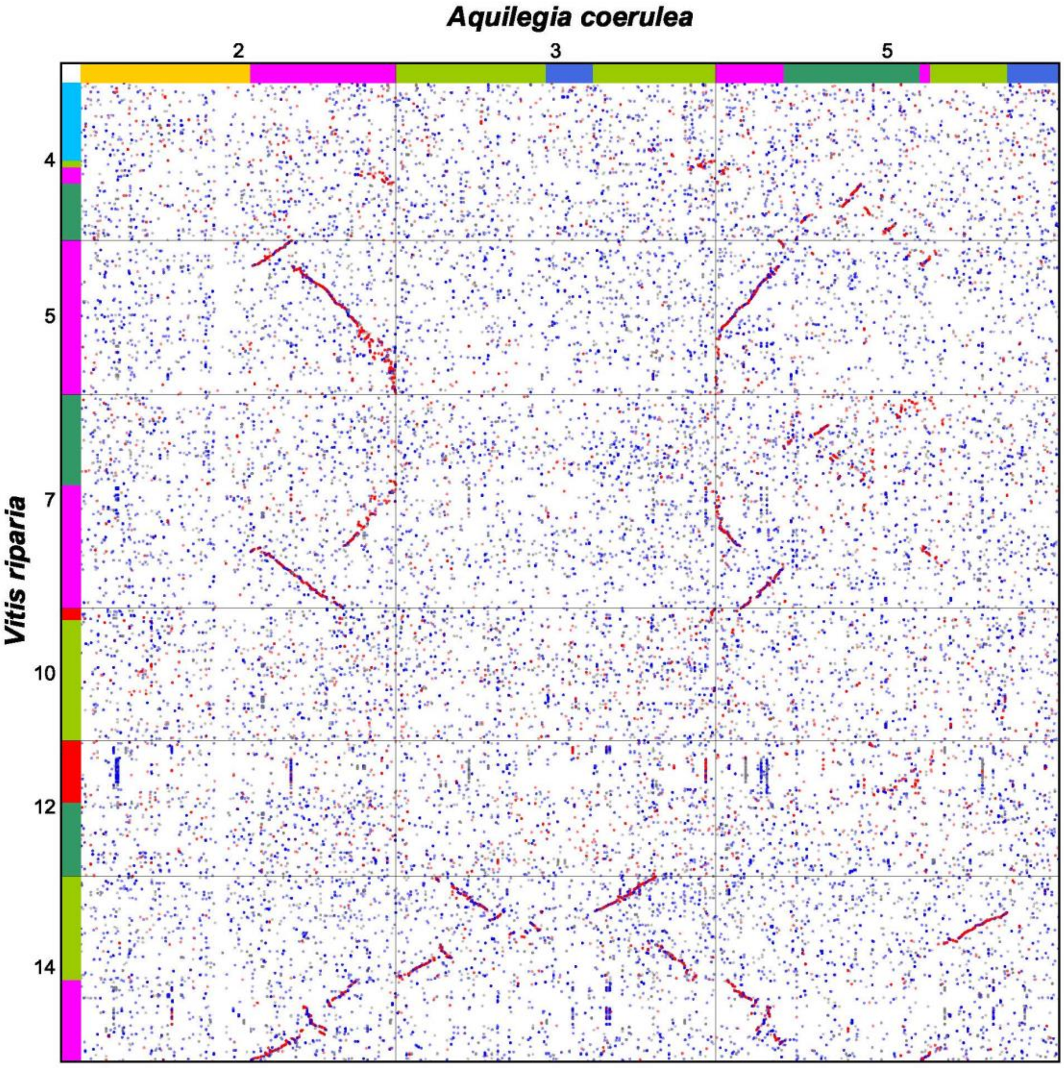


**Supplementary Fig. 7 Homologous dotplot between *Vitis riparia* and *Aquilegia coerulea.***

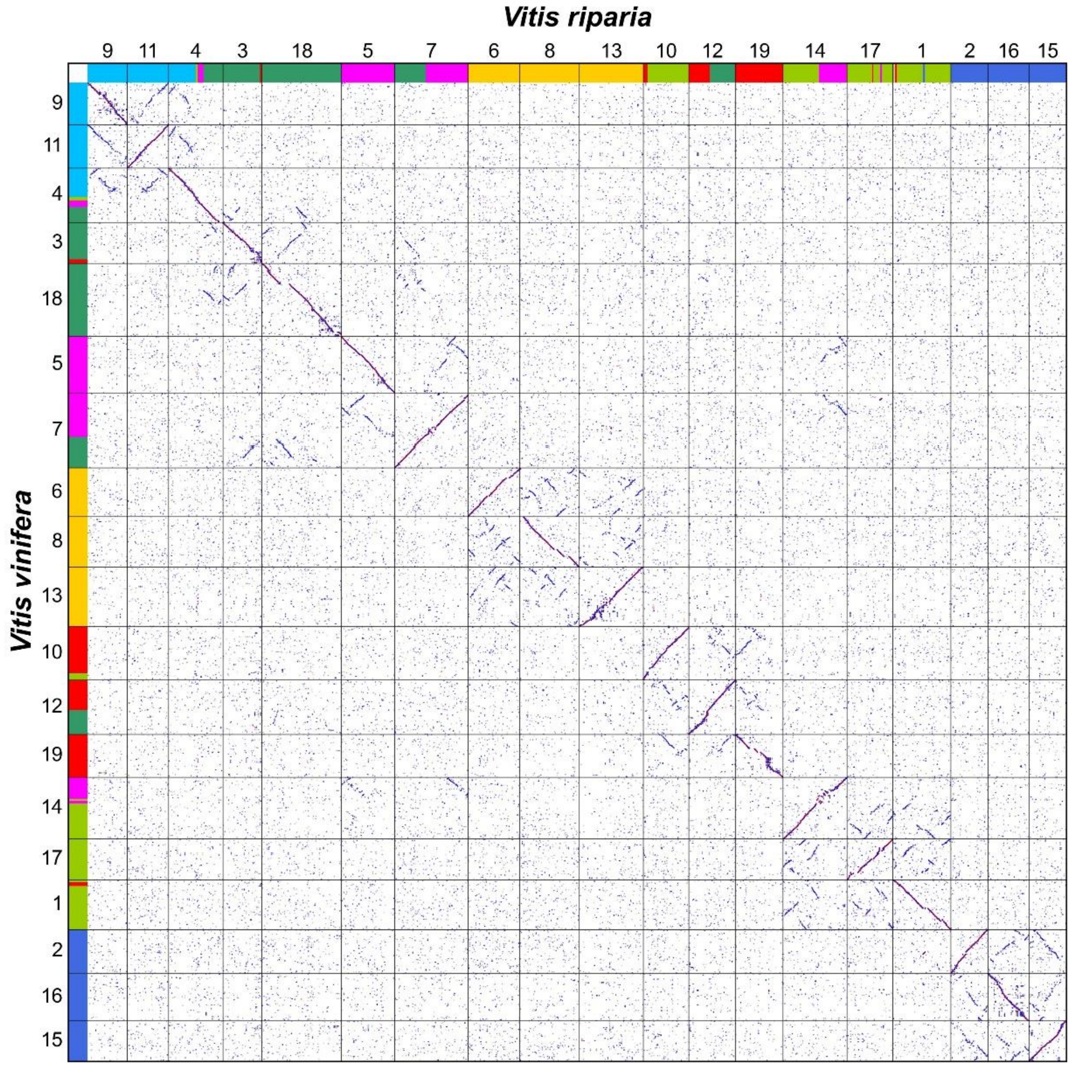


**Supplementary Fig. 8 Homologous dotplot between *Vitis vinifera* and *Vitis riparia.***
